# Reperfusion reshapes the temporal evolution of neurovascular injury after ischemic stroke

**DOI:** 10.64898/2026.09.03.744191

**Authors:** Maria Ardaya, Federico N. Soria, Aitzol García-Etxarri, Abraham Martín

**Affiliations:** Donostia International Physics Center (DIPC), San Sebastian, Spain; Achucarro Basque Center for Neuroscience, Leioa, Spain; Ikerbasque, Basque Foundation for Science, Bilbao, Spain

**Author notes:** Corresponding author: María Ardaya, Donostia International Physics Center (DIPC), Manuel Lardizabal Ibilbidea, 4, 20018 Donostia / San Sebastián, Gipuzkoa (Spain). Email address. Abraham Martín, Achucarro Basque Center for Neuroscience, Science Park of the UPV/EHU, 48940 Leioa, Spain.

**Keywords:** ischemia-reperfusion injury, *in vivo* two-photon imaging, vascular plasticity

## Abstract

Reperfusion is the cornerstone of acute ischemic stroke treatment, yet how it reshapes the temporal trajectory of neurovascular injury beyond the acute phase remains unclear. Here, we developed permanent (pStroke) and transient (tStroke) cortical ischemia mouse models compatible with longitudinal *in vivo* two-photon imaging to investigate the evolution of neurovascular injury following ischemia and reperfusion. Although reperfusion markedly reduced acute infarct development and neuronal loss, it failed to restore neurovascular homeostasis. Longitudinal two-photon imaging of ischemic mice revealed neuronal hyperexcitability and aberrant network synchronization, concomitant with impaired vascular remodeling and sustained vascular leakage following reperfusion. Histological analyses further demonstrated progressive neurodegeneration, chronic microglial activation and persistent alterations in the neurovascular unit despite improved preservation of brain tissue during the acute phase. Analysis of dextran permeability revealed size-selective BBB dysfunction, indicating that reperfusion induces prolonged impairment of vascular barrier properties rather than complete vascular recovery. Together, our findings demonstrate that reperfusion might reshape the temporal evolution of ischemic brain injury by limiting acute tissue damage while also promoting chronic neurovascular dysfunction, identifying persistent vascular instability as a potential therapeutic target for improving long-term recovery after ischemic stroke.

## INTRODUCTION

Ischemic stroke is one of the leading causes of mortality and long-term disability worldwide and results from the sudden interruption of cerebral blood flow, most commonly due to thrombotic or embolic arterial occlusion^1^. The resulting deprivation of oxygen and glucose initiates a complex cascade of metabolic failure, excitotoxicity, oxidative stress, and inflammation that rapidly culminates in neuronal death and tissue infarction^2^. Restoration of cerebral blood flow through intravenous thrombolysis with recombinant tissue plasminogen activator (rtPA) or mechanical thrombectomy remains the cornerstone of acute stroke treatment, substantially reducing infarct size and improving functional outcome when performed within an appropriate therapeutic window^3–6^. Although reperfusion effectively limits acute ischemic injury, increasing evidence indicates that restoration of blood flow can also initiate a secondary wave of tissue damage known as ischemia–reperfusion injury^7,8^. This process is characterized by oxidative stress, excitotoxicity, mitochondrial dysfunction, inflammatory activation and blood–brain barrier (BBB) disruption, all of which contribute to delayed neuronal degeneration, cerebral edema and hemorrhagic transformation^9–11^. BBB disruption is particularly consequential, as it promotes the infiltration of circulating plasma proteins and immune cells into the brain parenchyma, amplifying neuroinflammation and impairing tissue repair^11–13^. Despite these well-established mechanisms, it remains unclear how reperfusion reshapes the temporal evolution of neurovascular injury and why restoration of cerebral blood flow frequently fails to prevent chronic neurovascular dysfunction and long-term neurological impairment.

Progress in the study of the consequences of reperfusion has been limited by the lack of experimental models that simultaneously reproduce ischemia–reperfusion injury while allowing longitudinal visualization of neurovascular responses *in vivo*. The intraluminal middle cerebral artery occlusion (MCAO) model remains the gold standard for experimental stroke but typically produces extensive infarcts that limit detailed analysis of the ischemic penumbra and neurovascular unit^14,15^. Alternative approaches, including photothrombosis, electrocoagulation, endothelin-1 injection and magnetic nanoparticle-mediated occlusion, address specific experimental questions but generally induce permanent ischemia or provide limited access to the dynamic vascular events associated with reperfusion^16–19^. Consequently, the cellular and vascular mechanisms driving ischemia–reperfusion injury remain incompletely characterized.

To model primary ischemic damage and reperfusion-associated injury, here, we developed two complementary methods: permanent (pStroke) and transient (tStroke) focal cortical ischemia models in adult mice that are fully compatible with longitudinal *in vivo* two-photon imaging. This experimental platform enables real-time visualization of neuronal activity, vascular remodeling, BBB integrity and neuroinflammatory responses from the acute to the chronic stages of ischemic stroke, complemented by histological and behavioral analyses. Using these complementary models, we demonstrate that reperfusion reshapes the temporal evolution of ischemic brain injury by uncoupling acute neuronal preservation from chronic neurovascular dysfunction. Although reperfusion limits acute infarct development, it promotes persistent BBB disruption, progressive neurodegeneration and sustained neuroinflammation, identifying chronic vascular dysfunction as a central hallmark of ischemia–reperfusion injury. These findings provide new insight into the neurovascular mechanisms underlying long-term stroke progression and establish a versatile platform for investigating therapies aimed at preserving neurovascular integrity after reperfusion.

## METHODS

### Animals

Two-month-old male C57BL/6J mice (n = 242; body weight, 25 ± 2 g; Janvier Labs, France) were used in this study. All experimental procedures were approved by the Ethics Committee of the University of the Basque Country (UPV/EHU) under license M2024/007 and the corresponding local authorities. Animal care and experimental procedures were conducted in accordance with the European Directive 2010/63/EU on the protection of animals used for scientific purposes. This study is reported in compliance with the ARRIVE guidelines^20^.

### Experimental design

Three complementary experimental scenarios were designed to investigate the temporal evolution of ischemic injury following permanent (pStroke) or transient (tStroke) cortical ischemia.

The first scenario focused on the characterization of infarct development, neurodegeneration, neuronal activity, neurovascular unit alterations, glial reactivity and BBB disruption by histology, immunofluorescence and *in vivo* two-photon calcium imaging. To evaluate penumbra formation, mice were euthanized at 1, 3, 6 and 12 h after stroke induction, whereas brain atrophy and chronic pathological changes were assessed at 1, 3, 7, 15 and 30 days post-injury. A total of 92 mice were used for histological and immunofluorescence analyses, and 18 mice were used for longitudinal calcium imaging.

The second scenario investigated vascular remodeling and BBB integrity by longitudinal in vivo two-photon imaging. Vascular architecture was analyzed at 1, 3, 7, 15 and 30 days after stroke induction using 60 mice. An additional 24 mice were used to evaluate BBB size-selective permeability by intravascular administration of dextrans with different molecular weights.

The third scenario evaluated functional recovery using a battery of sensorimotor and cognitive behavioral tests performed at 1, 7, 15 and 30 days after stroke induction. A total of 48 mice were included in these behavioral analyses.

All experimental paradigms are summarized in the corresponding figures. Unless otherwise indicated, six animals per experimental group (sham, pStroke and tStroke) were included at each time point except at the time points assessed during the hours following ischemic onset to study penumbra progression, for which 4 animals were included per time point. For the acute penumbra analyses, four animals per group were used.

### Stroke models

#### Ligation of the somatosensory middle cerebral artery branch

Focal stroke models were induced via ligation of the middle cerebral artery branch which irrigate the right somatosensory cortex. Mice were anesthetized with 4% isofluorane for induction and 1.5% isofluorane for maintenance. Head hair was removed and skin sterilized with 70% alcohol. Somatosensory cortex was localized using a stereotaxic frame and a small craniotomy of about 1 mm was performed just above the branch. The branch structure is conserved in the same manner although with some inter-individual differences. The knot is performed just before the bifurcation of the somatosensory branch where the barrel cortex is located. Two approaches were developed: permanent stroke model (pStroke), where the perfusion is not restored, and transient stroke model (tStroke), where the perfusion is restored after 60 minutes of occlusion. After performing the model in both approaches, craniotomy was sealed with bone wax, skin sutured and sterilized with iodine povidone. The craniotomy was sealed with a glass coverslip and fixed to a metal head-plate for *in vivo* two-photon experiments. 0.1 mg/kg buprenorphine dose was injected for peri- and postoperative pain treatment. Animal’s welfare was supervised every day.

### Viral injection

Adeno-associated Syn.GCaMP6f.WPRE.SV40 virus (Addgene #100837-AAV9, US) was used to study calcium activity after stroke models. A virus volume of 400 nl (1×10¹³ vg/mL) was injected into layer II/III of the somatosensory cortex three weeks before performing the cranial window protocol and stroke models. A nanoinjector (WPI, #NANOLITER2020/300704 Injector, US) was used to inject the virus. In brief, animals were anesthetized with 4% isofluorane and placed in the stereotaxic frame. Anesthesia was maintained at 1.5% isofluorane for the whole experiment. Head hair was removed, skin sterilized with 70% alcohol and a skin incision was made to visualize skull bones. Injection coordinate was -1.0 AP, -3.0 ML and -0.3 DV in millimeters. A small craniotomy was made to introduce the glass pipette. After injection, the hole was sealed with bone wax, skin sutured and sterilized with iodine povidone. Cranial window was performed three weeks before viral injection. Peri- and postoperative pain management was conducted using subcutaneous injections of buprenorphine at a dose of 0.1 mg/kg. Mice were analyzed 24 and 72 hours after stroke models.

### Cranial window

Animals were anesthetized with 4% isoflurane and maintenance at 1.5% isoflurane during surgery. Mice were placed in a stereotaxic frame and somatosensory cortex (AP: -1; ML: -3; DV: -0.3 mm) was located. A 5-mm craniotomy was performed by slowly removing the skull with a drill (Foredom, US). Skull temperature was reduced with cold artificial cerebro-spinal fluid (aCSF) and air-puffed balloon during the process. Craniotomy was covered with a glass coverslip implant and glued with cyanoacrylate. aCSF was added between the brain surface and the glass implant. A metal plate was fixed to the skull with Superbond dental cement. This headplate allows head fixation to a holder in order to perform *in vivo* two-photon imaging with minimal head movement. Finally, skin was glued to the dental cement to avoid skin growth over the cranial window. Peri- and postoperative pain management was conducted using subcutaneous injections of buprenorphine at a dose of 0.1 mg/kg. Animals were supervised daily for each experiment.

### Two-photon imaging

A Femto-2D microscope (Femtonics, Hungary) equipped with a MaiTai Ti:Sapphire femtosecond Laser (Spectra-Physics, US) was used. Images from both set of experiments were registered at 1 Hz, 512x512 image size, 2 pixels of line average,1,37 μm/pixel, 525/50 and 620/60 filters and a water immersion 20× objective (NA 1.0, Olympus). Mice infected with Syn.GCaMP6f.WPRE.SV40 virus were used to study neuronal activity through calcium fluctuations. Mice were anesthetized with 3-4% isofluorane for induction and 0.5-1% isofluorane for maintenance during imaging experiments. After induction, mice were placed in the holder and kept under anesthesia. Two-photon laser was set to 920 nm and an intensity of 30 mW. Superficial neurons of layer II/III were analyzed in the sham group and in both stroke models. Stroke groups were analyzed at 24 and 72 hours. 53-56 neurons per conditions from four different animals (24 hours) and 20-24 neurons per condition from four different animals (72 hours).

For blood vessel analysis, 75 kDa dextran-TRITC was injected intravenously in the mouse tail vein using a 30G-insulin syringe just before imaging. Two-photon laser wavelength was set to 810 nm for TRITC and at 20 mW intensity. Superficial vessels were imaged from 0 to 200 µm in depth. Superficial middle cerebral artery branches and veins were analyzed. Extravasation experiment using two-photon imaging is described in *Extravasation assay* section.

### Two-photon image analysis

Calcium images were first motion-corrected to remove breathing artefacts. FIJI software (ImageJ)^21^ was used to downscale raw images to 256x256 pixels. Images were normalized to ΔF/F_0_, cell ROIs were manually traced and ΔF/F_0_ signal extracted for further analysis. ΔF/F_0_ values were processed in MATLAB and events thresholded to three standard deviations to extract raster plot data. ΔF/F_0_ traces were used to visualize neuronal activity in the different groups and peak frequency, peak amplitude and peak width (Hz) were extracted. Further analysis was performed using a MATLAB code developed by Jesús Pérez-Ortega and colleagues^22^. Raster data was analyzed to extract co-activity and similarity. Smooth filter of 1000 ms was used to compute z-score coactivity. Similarity values were extracted using Euclidean method and ward linkage method.

Maximal projections of 100 µm z-stack images were analyzed for vessel studies. Binary images of blood vessels were skeletonized for structural analysis^23,24^. Blood extravasation was measured with a surrounding mask of 30-µm band. Same approach was used for leakage of different size dextrans. FIJI software was used for analysis. Quantifications were manually performed except for vessel skeletonized and fractal dimension that were measured by Analyze Skeleton and Fractal count FIJI plugins^23,25^.

### Extravasation assay

*Evans blue injection:* 40 µl of Evans Blue (Merck #E2129, Germany) at 50 mg/ml was injected 60 minutes before mice euthanasia. The dye was injected intravenously in the mouse tail vein using a 30G-insulin syringe. Perfusion was later performed as described in *Histology and immunohistochemistry* section.

#### Injection of dextrans with different molecular weights

Mice were anesthetized with 4% isofluorane for induction and 1.5% isoflurane for maintenance. Mice were placed under the stereomicroscope, facing up and 1-cm skin was cut in the neck. Salivary glands were separated and the right carotid artery isolated. Fine bore polythene tubing (0.28mm ID; 0.61mm OD; Smiths Medical International, UK) was used for intracarotid administration. Tube pre-charged with the desire fluorescence dye was double-sutured to the carotid artery. Then, the skin was sutured and mice were placed under the two-photon objective using the head holder coupled to the anesthesia mouth cone. Three different molecular weight dextran were used to study vessel integrity after stroke models: 75 (Merck #T1162, Germany) and 40 kDa (Merck #53379, Germany). To combine them, 40 kDa dextran was conjugated with fluorescein (FITC) and 75 kDa dextrans was conjugated with TRITC. Bolus push was performed 2 minutes following the recording start. The length of the video was 30 minutes in total and the first 10 minutes were analyzed. Mice were kept under anesthesia during all the experiment. After 30 minutes of recording, mice were sacrificed as described in the following section.

### Histology and immunohistochemistry

Mice were anaesthetized with an intraperitoneal injection of pentobarbital (100 mg/kg), intracardially perfused using 4% paraformaldehyde in PBS and brain tissue overnight post-fixed. Fixed brains were sliced at 40 μm of thickness using the Leica VT 1200S vibratome (Leica Microsystems, Germany). Cresyl Violet staining was performed to assess infarct volume where tissue slices were incubated in the Cresyl Violet solution (Morphisto #11207.0025, Germany) and later dehydrated with an alcohol gradient. Xylol was used as a final step and slices were mounted with DPX mounting medium (Merck, Germany). Fluorojade-C staining was used to study neurodegeneration in the infarct area. Slides were incubated in 0.06% potassium permanganate solution for 15 minutes to reduce fluorescence background. After washing the potassium permanganate solution, slides were incubated in 0.0004% Fluorojade-C solution (Merck, #AG310, Germany) in 0.1% acetic acid for 30 minutes. Slides were kept drying overnight after a final washing step. The following day, slides were incubated in xylol before mounting them with DPX medium (Merck, Germany).

For immunofluorescence, tissue slices were permeabilized with 0.1% Triton and non-specific epitopes blocked with 1% Bovine Serum Albumin in PBS. Primary antibodies were incubated overnight and diluted in the same blocking solution. Guinea pig α-NeuN (Synaptic Systems #266004, Germany), chicken α-GFAP (Abcam #ab4674, UK), goat α-Iba1 (Abcam #ab5076, UK), guinea pig α-Aquaporin 4 (Synaptic Systems #429004, Germany), rabbit α-Collagen IV (Abcam #ab6586, UK), sheep α-Fibrinogen (Invitrogen #PA1-85429, US) and lectin (Merck #L0401, Germany) antibodies were used in this study. After primary antibody incubation, tissue was washed three times with PBS and secondary antibodies were added for one hour. Alexa fluor donkey α-goat 488, alexa fluor goat α-sheep 488, alexa fluor donkey α-chicken 546, alexa fluor donkey α-rabbit 555 and alexa fluor donkey α-guinea pig 594 were used as secondary antibodies (all from Thermo Fisher, US). Then, three new washing steps were conducted. Slices were mounted with Mowiol 4-88 Reagent l (Merck, Germany).

### Confocal microscopy acquisition and analysis

Confocal microscopy was performed using a Leica Stellaris (Leica Microsystems, Germany). Out of all slices collected in one dish, six coronal sections were chosen to represent the somatosensory cortex. Five regions were analyzed per slice, all located in the injured area. Series of z-stacks images were imaged (total z-depth of 8.4 µm) with a 40X (1.0 NA) oil immersion objective, zoom of 2, 1024x1024 image size and 0.14 μm/pixel. Double-positive expression of GCaMP6f and c-Fos was quantified to measure neuronal reactivity via the FIJI software (ImageJ)^21^. Fluorescence was visualized at 640 nm for Evans Blue quantification. Fluorescence profiles were analyzed using a custom ImageJ script (see https://github.com/SoriaFN/Tools). Double-positive expression of LEA/ColIV was used to count reactive vessels. AQP4 area and fragments were quantified using a 2 μm-band mask for LEA and ColIV^+^ vessels. The Analyze Skeleton plugin was used on the skeletonized segmented AQP4, to reveal the length and interconnectivity of the astrocyte end-feet of the vessel structure^23,25^. Cellular density was calculated as the number of cells/total cell number (% of total cell number). For NeuN, GFAP and Iba1, DAPI was used as a control and cells were considered positive when a band surrounded the nuclei exceeded a pre-established threshold for each marker.

### Behavioral tests

#### Open field test

We used this test to assess locomotor skills, analyzing total distance traveled. Mice were placed in the field for ten minutes before and after stroke induction. Training sessions were conducted for two weeks prior to the experimental tests.

#### Pole test

motor skills were also evaluated using the pole test, a widely used assay in animal models of ischemic stroke. Animals were trained prior to stroke induction, and scores were collected at 1, 7, 15, and 30 days post-stroke. Results represent the average of three trials conducted at 10-minute intervals per animal. Latency was defined as the time between task initiation and completion. Both prolonged latency (due to freezing) and shortened latency (due to premature falls) were considered indicative of motor deficits.

#### Cylinder Test

This assay evaluates bilateral motor involvement after MCAO. Animals were placed in a transparent glass cylinder (40 cm high, 20 cm wide), and the use of the forelimb on the affected side was monitored and documented. The first twenty contact attempts were analyzed. Animals from all experimental groups were assessed at 1, 7, 15 and 30 days after stroke models.

#### Adhesive removal test

This classical test was used to assess motor function by measuring the mouse’s ability to remove an adhesive tape attached to its paw. Animals underwent training sessions prior to testing and experimental evaluations were performed at 1, 7, 15, and 30 days post-stroke. Each session was recorded and latency was quantified both from task initiation to the first contact of the mouth with the adhesive (sensory function) and from task initiation to adhesive removal (motor dexterity).

#### Novel Object Recognition test

This assay evaluates short-term and long-term recognition memory in rodents by exploiting their innate tendency to explore unfamiliar items. This test was evaluated 30 days after stroke model induction. Animals underwent the familiarization session with two identical objects. Short-term memory performance was tested 3 hours after training with the familiar objects. Long-term memory was evaluated 24 hours after familiarization session. During each assessment, one of the original objects was substituted with a novel one. The discrimination index was calculated using the following formula: ([time exploring the novel object – time exploring the familiar object] / total exploration time).

### Statistical analysis

Data are expressed as means ± SD. Statistical analyses for two independent groups were conducted using an unpaired Student’s t-test. For comparisons involving more than two groups, one-way or two-way ANOVA, or mixed-effects models were applied, followed by Sidak’s post-hoc test to assess differences between stroke groups. Cumulative frequency distribution (CFD) was analyzed by Kolmogorov-Smirnov statistical test. Mann-Whitney U test was performed for 75 kDa dextran mask extravasation, pore size study and fluorescence profile of Iba1 and GFAP markers. The level of significance was set at p < 0.05, and statistical analyses were performed with GraphPad Prism version 8 software.

## RESULTS

### Reperfusion attenuates acute neuronal injury but promotes delayed neurodegeneration

Infarct volume and neurodegeneration were measured in the permanent (pStroke) and transient stroke (tStroke) models (Figure 1a). To study penumbra formation and brain atrophy, analyses were performed from the first hours after stroke onset through subsequent days (Figure 1b). Lesion volume was evaluated using Cresyl Violet staining (Figure 1c). In pStroke mice, infarct volume was almost fully established by 6 hours after stroke onset, whereas in tStroke mice it reached its maximum at 3 days (Figure 1d). Despite these differences in lesion development, brain atrophy progressed to a similar extent in both stroke models (Figure 1e). Neuronal degeneration was assessed using Fluoro-Jade C staining (Figure 1f). A temporal pattern similar to that observed for infarct volume was detected. pStroke mice exhibited significative neural degeneration as early as 6 hours after stroke onset, whereas tStroke mice showed peak neurodegeneration at 3 days (Figure 1g). However, at 30 days after stroke, tStroke mice displayed greater neurodegeneration than pStroke mice (Figure 1h), suggesting that reperfusion contributes to sustained neuronal death during the chronic phase of the disease. Despite these histopathological differences, only minor alterations were observed in sensorimotor and cognitive performance between the two stroke models at later stages (Supp. Figure 1). The only significative behavioral difference was observed in the novel object recognition test in which pStroke mice exhibited impaired short-term recognition memory 30 days after disease onset (Supp. Figure 1g). Overall, these findings indicate that the pStroke model induces earlier and more extensive irreversible brain damage than the transient model, whereas reperfusion is associated with enhanced long-term neuronal degeneration as well as a delayed evolution of the infarct.

**Figure 1.**
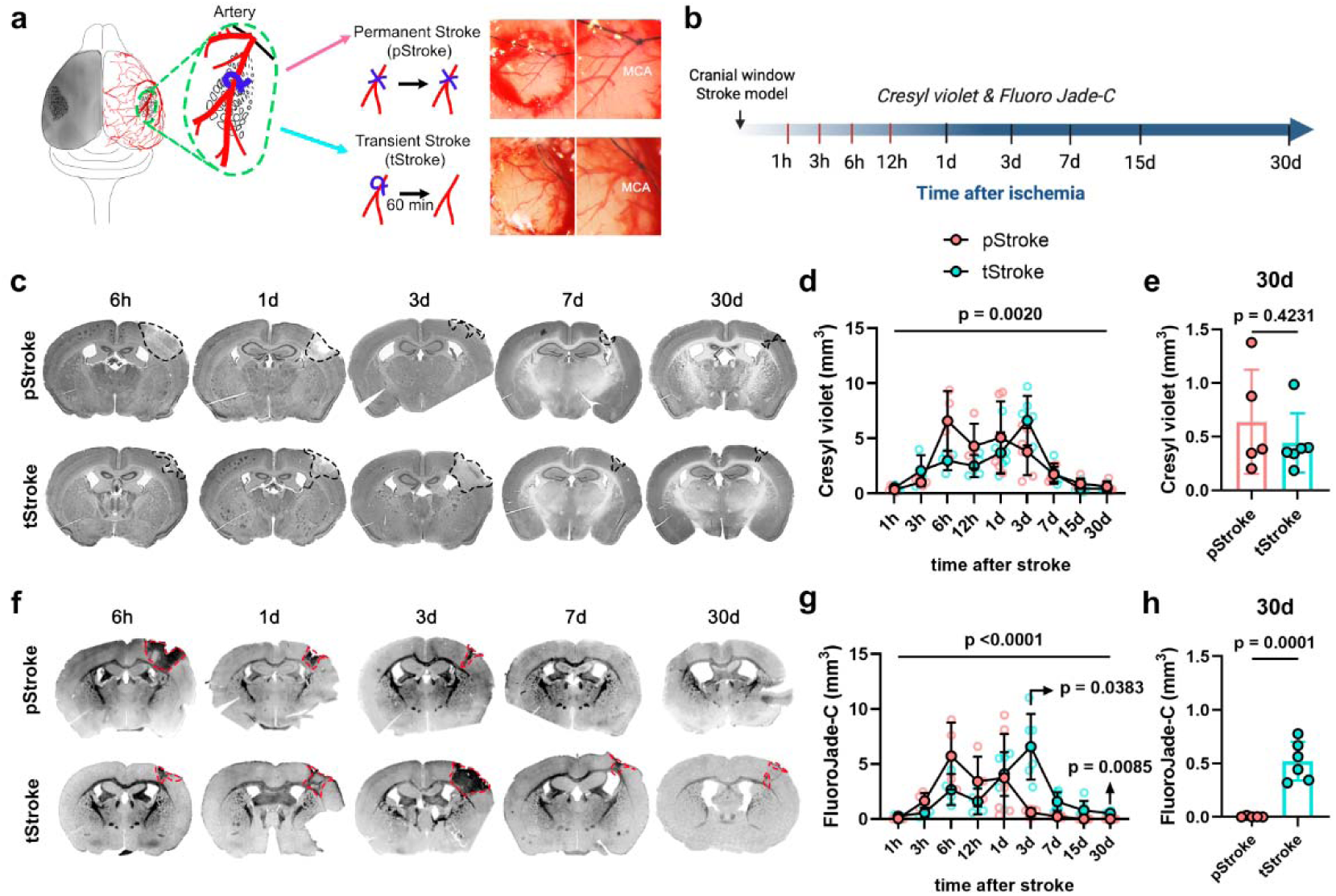
Reperfusion attenuates acute neuronal injury but promotes delayed neurodegeneration. Schematic representation of the permanent (pStroke) and transient (tStroke) focal cortical ischemia models in the right somatosensory cortex (a). Experimental design illustrating the time points analyzed during penumbra formation (hours) and brain atrophy (days) (b). Representative Cresyl Violet-stained brain sections from pStroke and tStroke mice at the indicated time points (c). Quantification of infarct volume revealed distinct temporal profiles, with a peak at 6 h in pStroke mice and at 3 days in tStroke mice (d). Despite these differences during the acute phase, both stroke models developed comparable brain atrophy 30 days after stroke onset (e). Representative FluoroJade-C staining showing neuronal degeneration in pStroke and tStroke mice over time (f). Quantification of FluoroJade-C-positive cells demonstrated an early peak of neurodegeneration in pStroke mice (6 h) and a delayed peak in tStroke mice (3 days) (g). Unlike infarct volume, neuronal degeneration remained elevated up to 30 days after reperfusion, indicating sustained delayed neuronal injury in tStroke mice (h). Data are presented as scatter dot plots showing individual animals with mean ± SD.

To further analyze neurodegeneration, neuron activity within the ischemic tissue was assessed by *in vivo* two-photon calcium imaging (Figure 2a). Animals were analyzed at 1 and 3 days after stroke onset (Figure 2b). An adeno-associated GCaMP6f virus was used to visualize intracellular calcium dynamics in neurons located in layer II/III of the right somatosensory cortex (Figure 2c). Raw fluorescence traces were extracted, concatenated, and normalized as ΔF/F₀ signals in sham, pStroke, and tStroke mice (Figure 2d). Quantification of GCaMP6f-positive neurons revealed a progressive reduction in neuronal number following stroke onset in both models (Figure 2e-f). Analysis of calcium transients showed that both stroke models exhibited increased firing frequency compared with sham animals at both 1 and 3 days after stroke, with no significant differences between pStroke and tStroke mice (Figure 2g). Peak amplitude showed a tendency to increase in both stroke models at 1 day, whereas a significant difference between stroke groups emerged at 3 days (Figure 2h). Moreover, peak width was significantly increased only in pStroke mice at 1 day after stroke (Figure 2i). Network analysis further revealed increased neuronal co-activity in both stroke models compared with sham animals at both time points, with significantly higher co-activity in pStroke than in tStroke mice (Figure 2j). Despite this increase in co-activity, peak similarity was reduced following stroke onset, and a significant difference between stroke groups was detected at 3 days (Figure 2k), suggesting that neuronal activity became increasingly aberrant and heterogeneous, consistent with pathological waves of depolarization rather than coordinated network activity. Together, these findings indicate that reperfusion attenuates acute neuronal loss while promoting persistent neuronal dysfunction, which is consistent with the delayed neurodegeneration observed at later stages of the disease.

**Figure 2.**
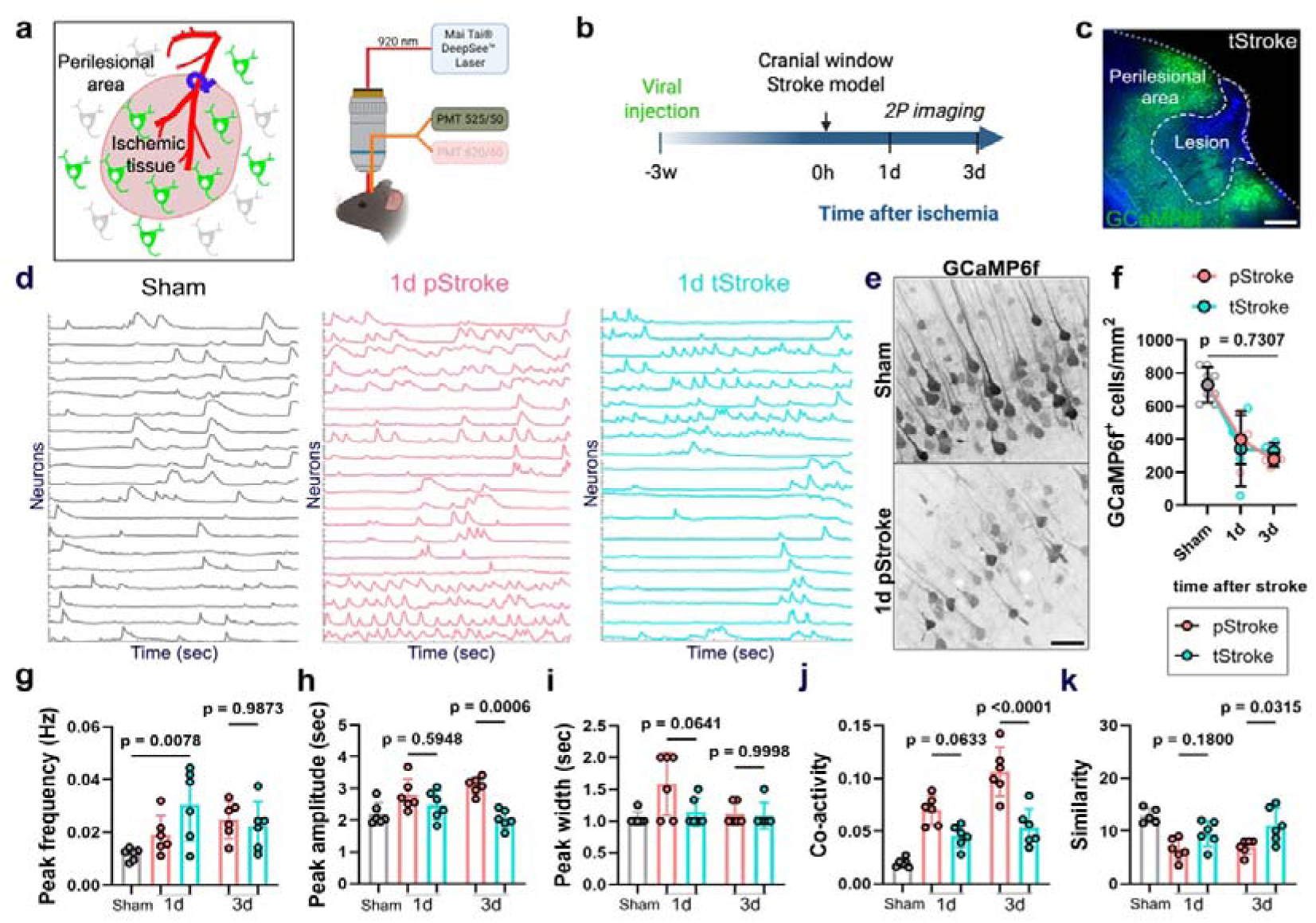
Reperfusion preserves neuronal viability but induces persistent alterations in neuronal activity. Schematic representation of adeno-associated virus (AAV)-mediated GCaMP6f expression in layer II/III neurons of the right somatosensory cortex and longitudinal in vivo two-photon calcium imaging (a). Experimental design showing calcium imaging performed at 1 and 3 days after stroke induction (b). Representative two-photon image of GCaMP6f-expressing neurons in the ischemic cortex (c). Representative normalized calcium traces (ΔF/F₀) recorded from sham, pStroke and tStroke mice (d). Representative images illustrating the progressive loss of GCaMP6f-positive neurons following stroke (e). Quantification of GCaMP6f-positive cells revealed a significant reduction in neuronal number in both stroke models at a and 3 days (f). Neuronal firing frequency increased in both pStroke and tStroke mice compared with sham controls at both time points (g). Peak amplitude showed a trend toward increased activity in both stroke models, with significant differences between pStroke and tStroke mice at 3 days (h). Peak width was selectively increased in pStroke mice at 1 day (i). Neuronal co-activity was significantly elevated after stroke, with higher synchronization observed in pStroke than in tStroke mice (j). In contrast, peak similarity was reduced in both stroke models, with significant differences between groups at 3 days, indicating the emergence of aberrant neuronal activity patterns following ischemia (k). Scale bars = 500 µm (c) and 50 µm (e). Data are presented as scatter dot plots showing individual animals with mean ± SD.

### Permanent ligation induces enhanced neurovascular plasticity

We next analyzed vascular remodeling following stroke models by *in vivo* two-photon imaging. The ligated artery and the adjacent vein were longitudinally analyzed in both stroke models (Figure 3a). To visualize the cerebral vasculature, 75-kDa tetramethylrhodamine (TRITC)-conjugated dextran was intravenously injected via the tail vein immediately before imaging (Figure 3b). Vascular imaging was performed at 1, 3, 7, 15 and 30 days after stroke onset (Figure 3c). The ligated artery was examined through the experimental period (Figure 3d) and a persistent intravascular clot was observed at 3 days only in pStroke mice (Supp. Figure 2a). Quantitative analysis revealed an increase in arterial vessel area following stroke onset in both models, whereas vessel diameter progressively decreased with a similar temporal profile (Figure 3e). Arterial vessel diameter decreased similarly in both stroke models (Figure 3e). In contrast, fractal dimension analysis showed that the arterial network exhibited greater structural complexity in pStroke than in tStroke mice (Figure 3e). Likewise, vessel density and tortuosity also differed between stroke models, whereas vessel segment partitioning remained unchanged (Supp. Figure 2b), suggesting that permanent ligation elicited a more pronounced vascular remodeling response. The vein adjacent to the ligated artery was also analyzed. Venous herniation was observed following stroke onset, suggesting increased local hemodynamic pressure (Figure 3f). Despite the methodological differences between the two stroke models, both groups exhibited comparable changes in venous area, diameter, and fractal dimension over time (Figure 3g). However, pStroke mice displayed greater venous density and tortuosity values compare to the tStroke mice (Supp. Figure 2c), whereas vessel segment partitioning remained unchanged (Supp. Figure 2c). These findings suggested that reperfusion attenuated vascular remodeling while inducing venous alterations comparable to those observed after permanent arterial occlusion.

**Figure 3.**
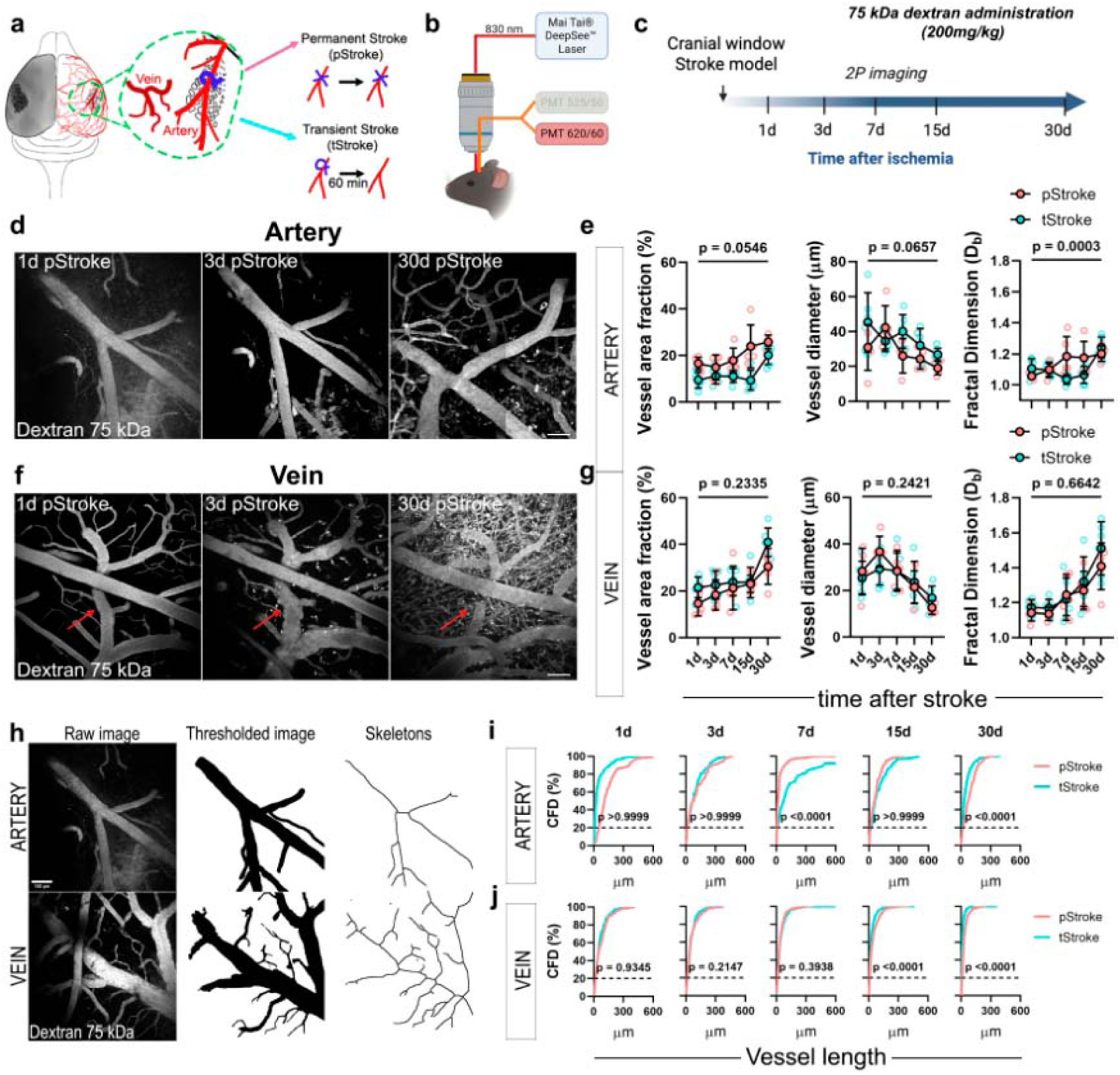
Reperfusion delays vascular remodeling after ischemic stroke. Schematic representation of the permanent (pStroke) and transient (tStroke) focal ischemia models (a). Experimental strategy for longitudinal in vivo two-photon imaging following intravenous administration of 75 kDa TRITC-dextran to visualize the cerebral vasculature (b). Experimental timeline (c). Representative two-photon images of the ligated artery in pStroke mice at 1, 3 and 30 days after stroke induction (d). Quantification of vessel area, vessel length density, vessel diameter and fractal dimension in the ligated artery of pStroke and tStroke mice (e). Representative images showing herniation of the vein adjacent to the ligated artery at 1, 3 and 30 days after stroke (f). Quantification of vessel area, vessel length density, vessel diameter and fractal dimension in the herniated vein of pStroke and tStroke mice (g). Representative vessel skeletons used for morphometric analysis (h). Quantification of vessel fragment length in the ligated artery at 1, 3, 7, 15 and 30 days after stroke (i). Quantification of vessel fragment length in the herniated vein at the corresponding time points (j). Scale bars = 100 µm (d, f and h). Data are presented as scatter dot plots showing individual animals with mean ± SD.

To further characterize vascular architecture, skeleton analysis was performed on the ligated artery and the adjacent herniated vein (Figure 3h). In the arterial network, vessel fragment length vessel fragment lengths differed between stroke groups at 7 and 30 days (Figure 3i). In contrast, vein skeletons analysis revealed comparable fragment lengths in both stroke groups, although significant temporal changes were observed at 15 and 30 days after stroke (Figure 3j). These findings indicates that reperfusion delayed vascular remodeling and limits structural vascular plasticity within the ischemic territory.

### Prolonged blood leakage following reperfusion

BBB disruption is a hallmark of ischemia–reperfusion injury and contributes to stroke progression. To characterize BBB integrity over time, Evans Blue dye was intravenously administered at early time points of stroke onset (1, 3, 6 and 12 hours) to assess penumbra evolution and at later time points (1, 3, 7, 15 and 30) to evaluate BBB alterations during brain atrophy (Figure 4a). Evans blue extravasation was imaged and later quantified by immunofluorescence (Figure 4b), with macroscopic leakage also readily detectable by gross examination (Figure 4c). Distinct temporal patterns of BBB disruption were observed between pStroke and tStroke mice (Figure 4d). Consistent with the temporal profiles of infarct formation and neuronal degeneration revealed by Cresyl Violet and Fluoro-Jade C staining, pStroke mice exhibited an early peak of Evans Blue extravasation at 6 hours and 1 day after stroke onset, whereas tStroke mice displayed a delayed but more pronounced peak at 3 days (Figure 4e). Moreover, Evans Blue extravasation remained detectable in tStroke mice at 15 and 30 days, despite being nearly absent in pStroke mice (Figure 4f-g, respectively), suggesting that reperfusion promoted persistent BBB disruption at later stages of the disease.

**Figure 4.**
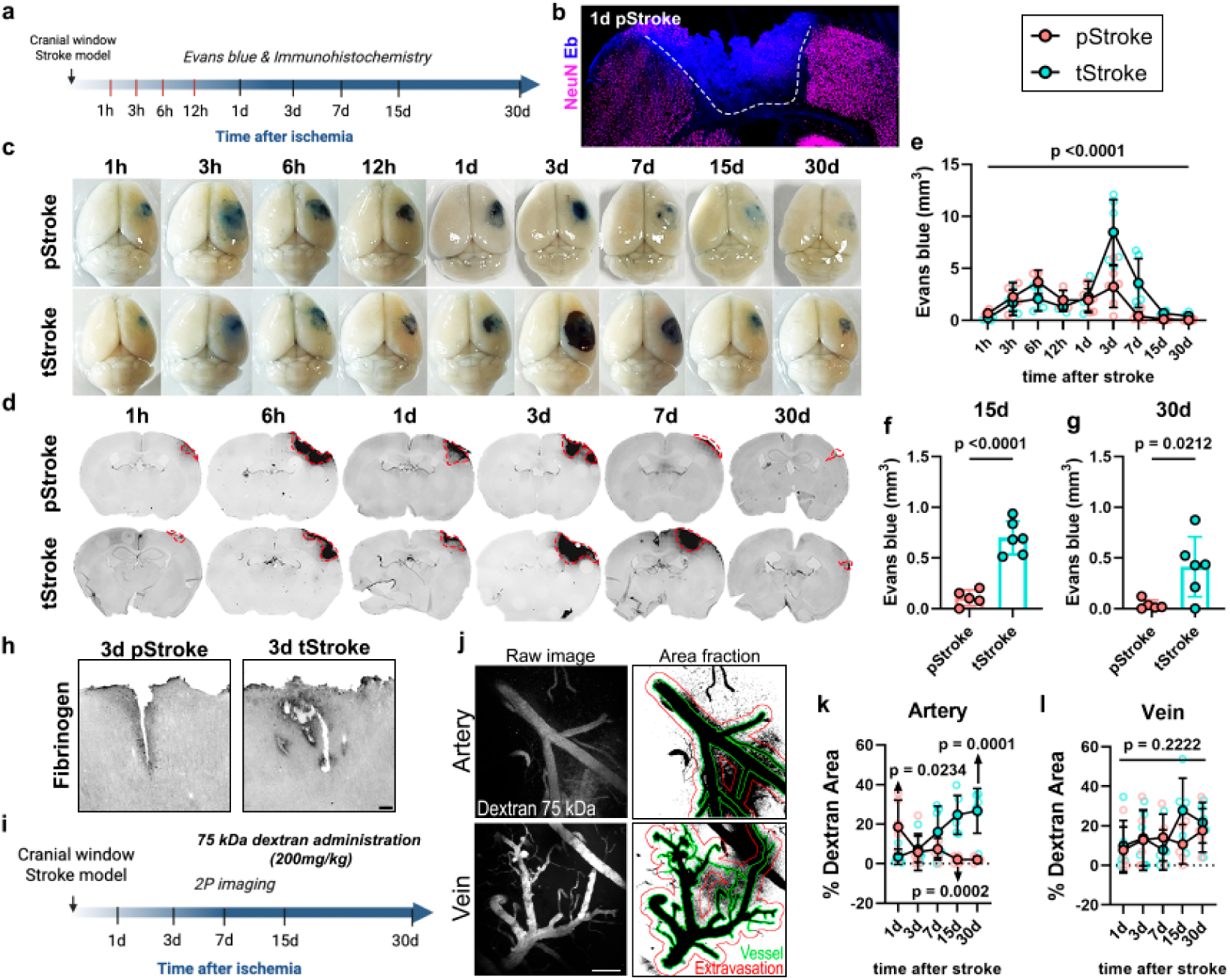
Reperfusion induces persistent blood–brain barrier disruption and chronic vascular leakage. Experimental design for the assessment of blood–brain barrier (BBB) permeability during penumbra formation (hours) and brain atrophy (days) following stroke induction (a). Evans Blue fluorescence was imaged in the ischemic hemisphere using infrared imaging (b). Representative images of Evans Blue extravasation in pStroke and tStroke mice at the indicated time points, visualized by gross inspection (c) and immunofluorescence (d). Quantification of Evans Blue extravasation revealed an early peak at 6 and 24 h in pStroke mice, whereas tStroke mice exhibited a delayed peak at 3 days after stroke (e). BBB leakage remained significantly elevated in tStroke mice at 15 (f) and 30 days (g), indicating persistent vascular dysfunction after reperfusion. Representative fibrinogen immunofluorescence images showing vascular leakage at 3 days after stroke in both models (h). Experimental strategy for longitudinal in vivo two-photon imaging of vascular leakage in a 30-µm region surrounding the ligated artery and the adjacent herniated vein (i). Following intravenous administration of 75 kDa TRITC-dextran, fluorescence images were thresholded to quantify dextran extravasation (j). Quantification of arterial leakage demonstrated greater early extravasation in pStroke mice that progressively declined, whereas leakage increased over time in tStroke mice (k). No significant differences in dextran extravasation were detected in the herniated vein between stroke models (l). Scale bars = 500 µm (b) and 100 µm (h–j). Data are presented as scatter dot plots showing individual animals with mean ± SD.

Vascular injury was further evaluated by fibrinogen immunofluorescence, which revealed leakage from large blood vessels and vascular reactivity in both stroke models (Figure 4h). BBB integrity was further analyzed by *in vivo* two-photon imaging (Figure 4i). The ligated artery and the adjacent herniated vein were visualized following intravenous injection of 75 kDa TRITC-conjugated dextran (Figure 4j). Consistent with the Evans Blue analyses, blood extravasation around the ligated artery was greater in pStroke mice at day 1, while reperfusion showed persistent vascular leakage at 7 days onwards, during the chronic phase of the disease (Figure 4k).

In contrast, no significant differences in blood extravasation were detected between stroke models in the adjacent vein (Figure 4l). Together, these findings suggest that although tStroke animals displayed smaller infarct volumes, reperfusion induced a chronic alteration of the vascular integrity and sustained BBB dysfunction.

To further characterize BBB permeability, fluorescent dextrans of different molecular weights were used to assess size-dependent vascular permeability by in vivo two- photon imaging (Figure 5a). vascular permeability was assessed following intracarotid administration of 75- and 40-KDa dextrans, and the ligated artery and adjacent herniated vein were analyzed at 1 and 3 days after stroke onset in both models (Figure 5b). 75 kDa dextran extravasation was detected in the ligated artery of pStroke mice at day 1 after stroke, with no further increase at 3 days (Figure 5c). In contrast, extravasation of 40 kDa dextran was significantly increased in tStroke mice at 1 day (Figure 5d) suggesting that both stroke models exhibited distinct temporal and size- selective patterns of BBB disruption. Despite these findings, no significant differences between stroke models were observed in the herniated vein for either 75 kDa (Figure 5e) or 40 kDa (Figure 5f) dextran at either time point. Together, these findings suggest that reperfusion limits the extravasation of high-molecular-weight tracers while promoting persistent permeability to smaller molecules, consistent with sustained low- pore BBB dysfunction following transient ischemia.

**Figure 5.**
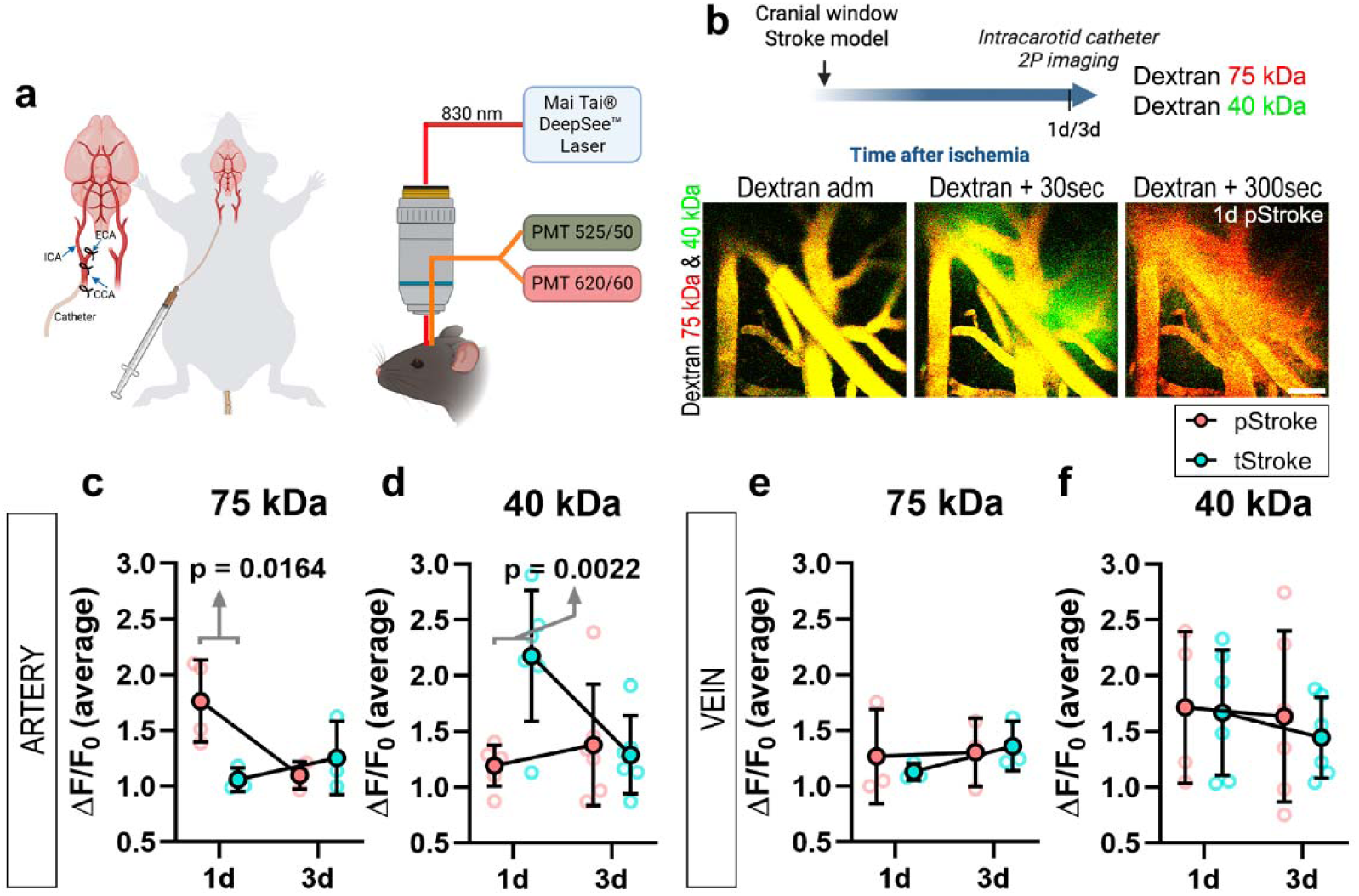
Reperfusion alters the size-selective permeability of the blood–brain barrier. Schematic representation of the intracarotid dextran administration protocol and in vivo two-photon imaging setup used to assess blood–brain barrier (BBB) size- selective permeability (a). Experimental timeline and representative two-photon images showing the distribution of 75 kDa (red) and 40 kDa (green) dextrans at baseline, 30 s and 300 s after injection (b). Quantification of dextran extravasation in the ligated artery revealed greater leakage of the 75 kDa dextran in pStroke mice than in tStroke mice at 24 h after stroke induction, indicating the presence of larger vascular pores during the acute phase (c). In contrast, 40 kDa dextran extravasation was greater in tStroke mice than in pStroke mice at 3 days, consistent with sustained permeability to smaller circulating molecules following reperfusion (d). No significant differences between stroke models were observed in the herniated vein for either the 75 kDa (e) or the 40 kDa (f) dextran. Scale bar = 100 µm (b). Data are presented as scatter dot plots showing individual animals with mean ± SD.

### Permanent occlusion induced an early vascular response involving basal lamina and astrocytic-end-feet

We next analyzed key components of the neurovascular unit during the acute phase of penumbra formation (hours) and the subsequent development of brain atrophy (days) (Figure 6a). Vascular compartmentalization was assessed by immunofluorescence of Lectin (LEA) to label endothelial cells, Collagen IV (ColIV) to identify the basal lamina and Aquaporin 4 (AQP4) as a marker of astrocytic end-feet (Figure 6b). We first analyzed the expression of ColIV in LEA-positive vessels following permanent and transient stroke models (Figure 6c). Expression of ColIV increased after stroke onset with a higher peak at 6 and 12 hours in pStroke mice (Figure 6d). Its overexpression was mainly localized in the core of the lesion (Figure 6e). Fractal dimension analysis further revealed differences in the structural complexity of ColIV-positive vascular fragments between stroke models (Figure 6f). Likewise, ColIV fragments size differed between pStroke and tStroke mice, particularly at 3 and 6 hours after stroke onset (Figure 6g) suggesting that basal lamina remodeling differs according to the presence or absence of reperfusion.

**Figure 6.**
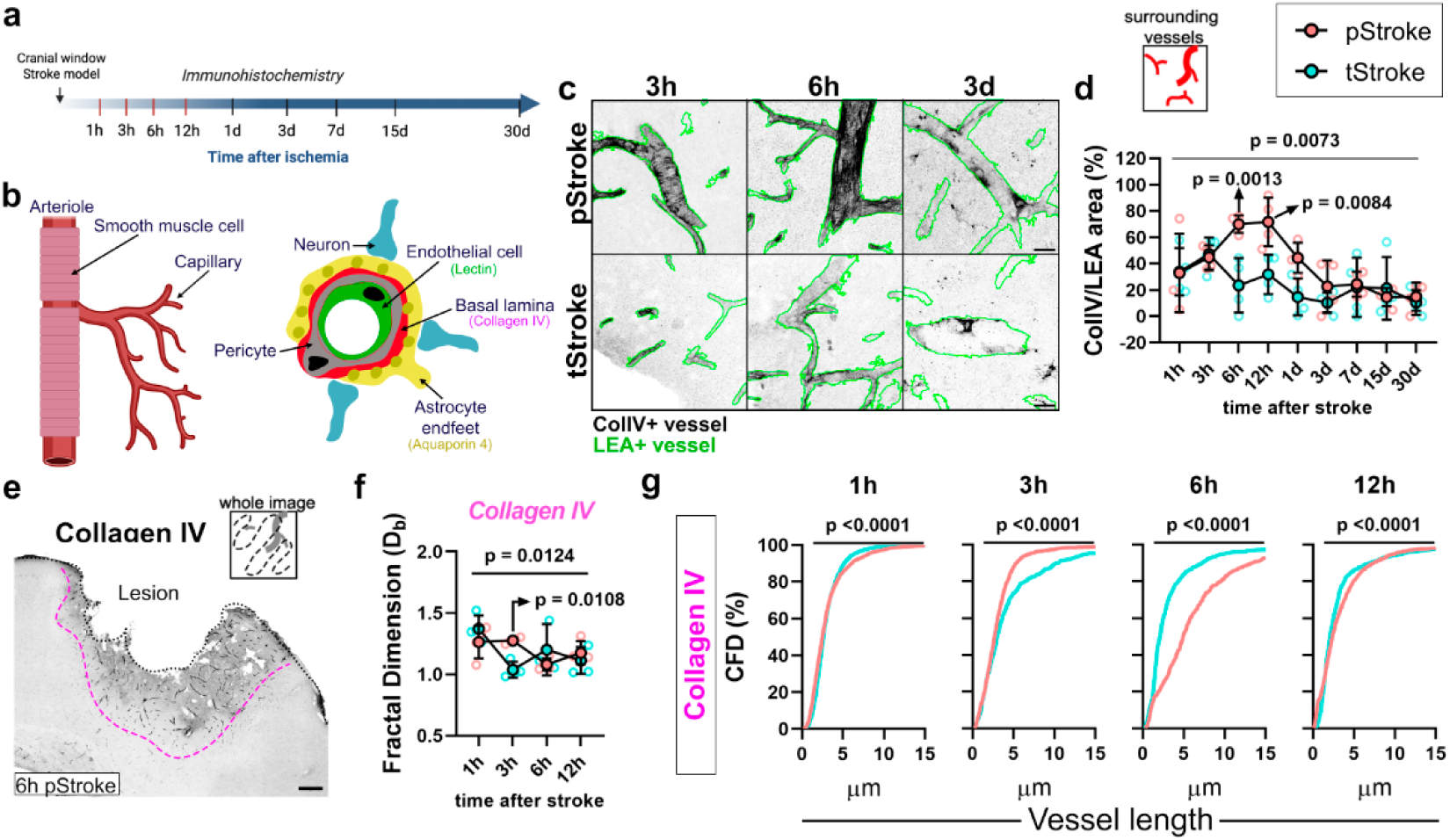
Reperfusion attenuates the early neurovascular unit response after ischemic stroke. Experimental timeline for the analysis of neurovascular unit components during penumbra formation (hours) and brain atrophy (days) following stroke induction (a). Schematic representation of the vascular compartments analyzed, including endothelial cells (LEA), the vascular basement membrane (collagen IV, ColIV) and astrocytic endfeet (AQP4) (b). Representative images of ColIV immunofluorescence (black) and LEA-positive vessels (green) at 3 h, 6 h and 3 days after stroke induction (c). Quantification of ColIV expression in LEA-positive vessels revealed a transient increase at 6 and 12 h exclusively in pStroke mice (d). Representative low-magnification image illustrating ColIV expression in the ischemic cortex of a pStroke mouse at 6 h (e). Fractal dimension analysis demonstrated significant differences in ColIV structural complexity between pStroke and tStroke mice (f). Quantification of ColIV fragment size revealed significant differences between stroke models during the acute phase (1, 3, 6 and 12 h), indicating distinct patterns of vascular basement membrane remodeling (g). Scale bars= 20 µm (c) and 200 µm (e). Data are presented as scatter dot plots showing individual animals with mean ± SD.

We next measured AQP4 expression by immunofluorescence (Supp. Figure 3a). We observed accumulation of AQP4 in ColIV-positive vessels (Supp. Figure 3b) suggesting a close spatial association between astrocytic end-feet and the vascular basal membrane following stroke. AQP4 expression was significantly increased in pStroke mice at 6 hours compare with tStroke mice (Supp. Figure 3c). Moreover, a positive correlation between ColIV and AQP4 expression was observed only in pStroke mice hours after the onset (Supp. Figure 3d) suggesting a strong relation between these vascular elements in the critical ischemic temporal window. Low-magnification imaging further revealed marked changes in AQP4 distribution between the healthy and ischemic tissue (Supp. Figure 3e). Although no significant differences in AQP4 fragment complexity was detected between stroke groups (Supp. Figure 3f), AQP4 fragments size changed over time (Supp. Figure 3g), with a greater proportion of large AQP4-positive structures observed in pStroke mice at 6 hours (Supp. Figure 3h). Together, these findings demonstrate coordinated remodeling of the vascular basal membrane and astrocytic end-feet during the early phase of ischemic injury, with more pronounced alterations following permanent arterial occlusion than after reperfusion.

### Reperfusion is associated with early glial activation and sustained chronic neuroinflammation

We next evaluated microglia/macrophage and astrocyte expression following permanent and transient stroke at multiple time points after disease onset to study glial response (Figure 7a). Glial reactivity was assessed by immunofluorescence in both the lesion core and the perilesional area (Figure 7b). At day 1, Iba1-positive cells were predominantly localized within the lesion core in pStroke mice, whereas they were mainly distributed throughout the perilesional region in tStroke mice (Figure 7c). Quantification revealed that Iba1 expression peaked at 7 days in pStroke mice, while tStroke mice exhibited an earlier peak at 3 days within the lesion core, which remained elevated at 7, 15, and 30 days after stroke onset (Figure 7d). GFAP expression followed a similar temporal profile in the lesion core, although prolonged astrocyte reactivity during the chronic phase was not observed in tStroke mice (Figure 7e). The total number of Iba1-positive cells was consistently lower in the perilesional area than the core of the lesion (Figure 7f). However, GFAP-positive cells occupied a larger area in the perilesional area, reaching maximal levels at 7 days in pStroke mice and at 3 days in tStroke mice (Figure 7g). These findings suggest that reperfusion accelerates the early glial response while promoting prolonged microglial activation during the chronic phase of ischemic injury.

**Figure 7.**
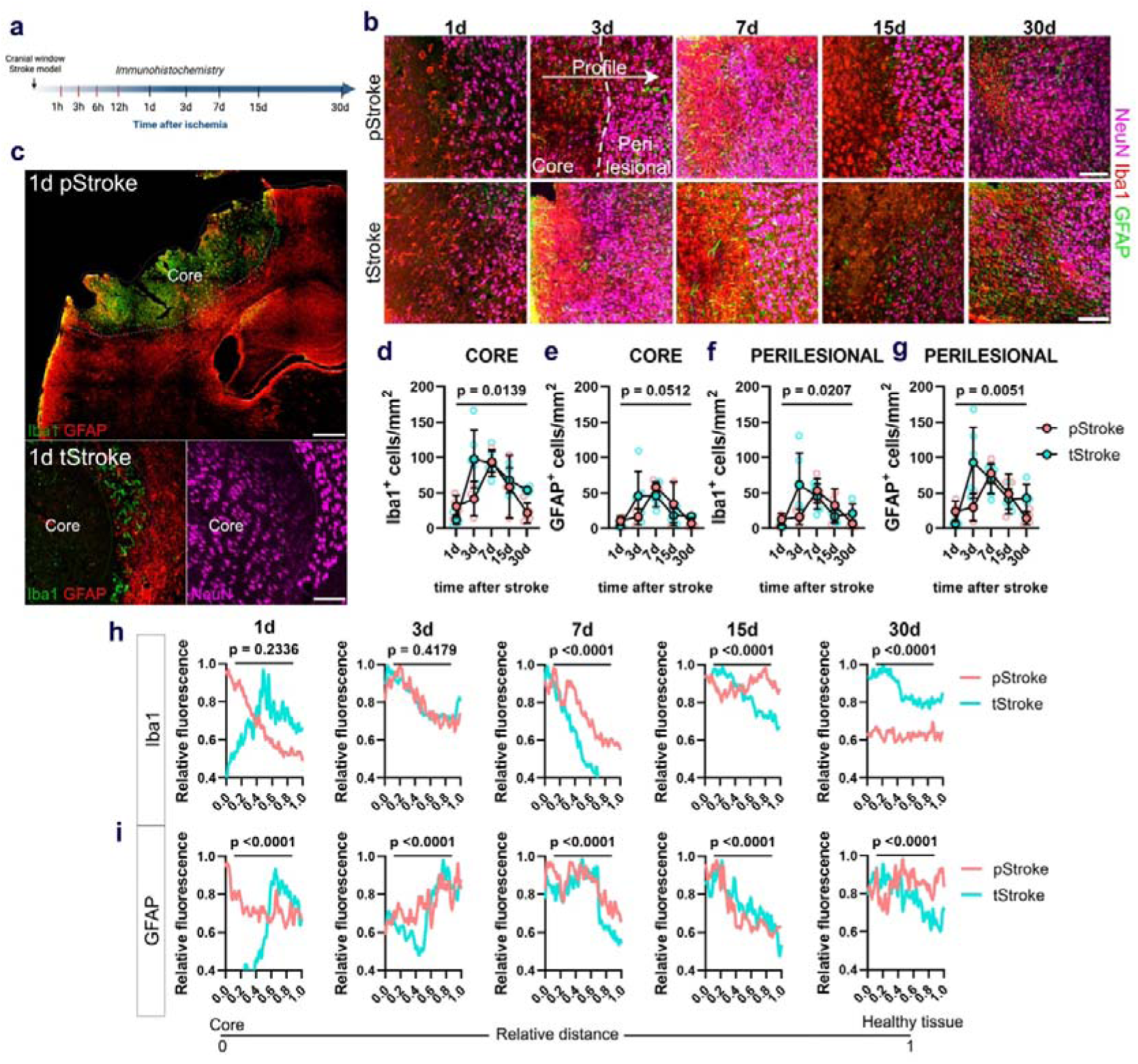
Reperfusion promotes persistent glial activation following ischemic stroke. Experimental timeline for the analysis of glial reactivity after permanent (pStroke) and transient (tStroke) focal ischemia (a). Representative immunofluorescence images of Iba1-positive microglia/macrophages and GFAP- positive astrocytes in pStroke (top) and tStroke (bottom) mice at 1, 3, 7, 15 and 30 days after stroke induction (b). Representative images illustrating the spatial distribution of Iba1-positive cells at 24 h, showing preferential localization within the lesion core in pStroke mice and in the peri-lesional region in tStroke mice (c). Quantification of Iba1 immunoreactivity in the lesion core revealed a peak at 7 days in pStroke mice, whereas tStroke mice exhibited an earlier peak at 3 days that remained elevated throughout the chronic phase (d). Quantification of GFAP immunoreactivity in the lesion core showed a similar temporal profile but without sustained chronic activation in tStroke mice (e). Quantification of Iba1-positive cells in the peri-lesional area showed no significant differences between stroke models (f). In contrast, GFAP- positive cells increased in the peri-lesional region, peaking at 3 days in tStroke mice and at 7 days in pStroke mice (g). Fluorescence intensity profiles of Iba1 (h) and GFAP (i) were generated from the lesion core toward the surrounding healthy tissue to evaluate the spatial distribution of glial reactivity at 1, 3, 7, 15 and 30 days after stroke induction. Scale bars = 100 µm (b), 500 µm and 100 µm (c). Data are presented as scatter dot plots showing individual animals with mean ± SD.

To further characterize the spatial distribution of glial activation, fluorescence intensity profiles for Iba1 and GFAP were generated along a transect extending from the lesion core to the surrounding healthy tissue at 1, 3, 7, 15, and 30 days after stroke onset (Figure 7h and i, respectively). Iba1 fluorescence profiles differed significantly between stroke models during the chronic phase, with sustained signal intensity in the lesion core of tStroke mice (Figure 7h), consistent with persistent microglial activation following reperfusion. The preferential localization of Iba1-positive cells within the perilesional region of tStroke mice at 1 day was also evident from the fluorescence profiles, although these differences did not reach statistical significance (Figure 7h). In contrast, GFAP fluorescence profiles differed significantly between the two stroke models at all analyzed time points (Figure 7i), indicating distinct astrocytic responses depending on the type of ischemic injury. These findings suggest that both immune cell activation and astrocyte reactivity exhibit differential temporal profiles according to the stroke signature.

## DISCUSSION

This study demonstrates that reperfusion fundamentally reshapes the temporal evolution of ischemic brain injury and subsequent functional recovery. Although restoration of cerebral blood flow effectively limits acute infarct development and neuronal loss, it also initiates a distinct phase of neurovascular pathology characterized by persistent BBB dysfunction, progressive neurodegeneration, and sustained neuroinflammation. These observations suggest that ischemia–reperfusion injury is a major determinant of the long-term neurological consequences of stroke and highlight chronic vascular dysfunction as a key pathological process that persists despite successful recanalization.

Within this framework, the permanent occlusion model (pStroke) primarily reproduced the acute consequences of primary ischemic injury, characterized by rapid anoxic– ischemic cell death, whereas the transient occlusion model (tStroke) more closely recapitulated the delayed pathological processes associated with ischemia–reperfusion injury. Rather than representing interchangeable stroke models, pStroke and tStroke capture distinct temporal trajectories of ischemic injury, providing complementary experimental platforms to investigate primary ischemic damage and chronic reperfusion-induced pathology.

Consistent with previous studies using permanent occlusion models, including the Bengal Rose photothrombotic model, permanent ischemia induced larger infarcts and more extensive neurodegeneration during the acute phase^16,26^. In contrast, transient ischemia reduced acute tissue damage but promoted delayed neuronal degeneration, in agreement with previous reports describing the detrimental long-term consequences of reperfusion injury^27,28^, suggesting that blood recanalization after ischemic stroke may have damaging long-term effects. Although thrombolysis and mechanical thrombectomy remain the most effective therapeutic strategies for limiting infarct expansion^29,30^, our findings suggest that restoration of blood flow does not terminate the pathological process but instead redirects it toward a chronic neurovascular response.

Two-photon calcium imaging further demonstrated that both stroke models developed early neuronal hyperactivity, consistent with previous observations^26,31–34^. Increased neuronal excitability has been proposed to promote the recruitment and activation of peripheral immune cells, thereby amplifying neuroinflammatory responses^35–37^. Our findings extend these observations by showing that, despite preserving a larger population of neurons during the acute phase, reperfusion induces persistent abnormalities in neuronal activity that may contribute to delayed neuronal degeneration. These data suggest that acute neuronal survival and long-term neuronal integrity are not necessarily coupled following successful reperfusion.

This dissociation between acute preservation and chronic dysfunction extended beyond neurons to the vascular compartment. Reperfusion failed to restore neurovascular homeostasis. Permanent ischemia induced a more robust vascular remodeling response, whereas reperfusion resulted in persistent vascular dysfunction despite smaller infarct volumes. Longitudinal two-photon imaging revealed impaired arterial remodeling together with sustained alterations in the adjacent venous compartment, suggesting that restoration of blood flow modifies the normal vascular repair program. Previous studies have demonstrated that vascular remodeling contributes to tissue repair after ischemic stroke^13,38,39^, whereas increasing evidence also supports an active contribution of the cerebral venous system to stroke pathology^40^. Our findings reinforce this concept by demonstrating that chronic venous abnormalities accompany persistent vascular dysfunction after reperfusion.

This impaired vascular remodelling was accompanied by persistent disruption of BBB integrity, suggesting that the failure to restore normal vascular architecture has direct consequences for barrier function. Evans Blue extravasation demonstrated that BBB disruption followed distinct temporal profiles in both stroke models, with transient ischemia exhibiting prolonged vascular leakage during the chronic phase. In vivo two- photon imaging further revealed dynamic BBB disruption within individual vessels, whereas dextran permeability assays demonstrated size-selective vascular leakage, indicating that reperfusion does not simply increase vascular permeability but alters the selective barrier properties of the BBB. This prolonged vascular instability may represent a central mechanism linking vascular injury to chronic neurodegeneration, as sustained leakage facilitates continuous exposure of the brain parenchyma to circulating plasma proteins, inflammatory mediators and peripheral immune cells, thereby maintaining a pro-inflammatory microenvironment that impairs tissue repair^34–37^. The analyses of collagen IV and AQP4 further support the existence of distinct neurovascular responses following permanent ischemia and reperfusion. Surprisingly, collagen IV expression increased rapidly after permanent ischemia, in contrast to previous reports describing basal membrane degradation during ischemia–reperfusion injury^13,38^. This early accumulation may reflect transient remodeling of the vascular basal membrane before proteolytic degradation or differences in the temporal dynamics between permanent and transient ischemia. Interestingly, early collagen IV accumulation was associated with greater vascular remodeling during the chronic phase, suggesting that basal membrane remodeling may contribute to vascular repair. Likewise, AQP4 expression increased rapidly after permanent ischemia and closely correlated with collagen IV expression, indicating coordinated remodeling of astrocytic end-feet and the vascular basal membrane. Although AQP4 overexpression has traditionally been associated with cytotoxic edema, studies using AQP4-deficient mice suggest a biphasic role in ischemic stroke, being detrimental during the acute phase but beneficial for edema resolution and tissue repair during recovery^39,40^. Together, these findings suggest that efficient remodeling of the neurovascular unit may represent an adaptive response that facilitates vascular stabilization after ischemic injury.

Finally, our data indicate that reperfusion profoundly influences the inflammatory response. Although glial activation is essential for tissue repair, transient ischemia induced sustained microglial activation that persisted throughout the chronic phase, whereas astrocytic responses followed distinct temporal dynamics. These observations suggest that prolonged BBB dysfunction may impair the resolution of neuroinflammation by maintaining continuous exposure of the injured brain to circulating inflammatory signals. Previous studies have demonstrated that local microglia play a central role in coordinating post-stroke inflammation and tissue repair^34–37^, supporting the hypothesis that therapeutic strategies aimed at preserving BBB integrity while modulating peripheral immune cell infiltration may improve long-term neurological recovery.

Together, our findings support a conceptual shift in ischemic stroke pathophysiology. Rather than representing the end of ischemic injury, successful reperfusion initiates a prolonged phase of neurovascular dysfunction characterized by persistent BBB disruption, chronic neuroinflammation and progressive neurodegeneration. These results identify vascular protection after reperfusion as a major therapeutic challenge and suggest that preserving neurovascular integrity may be as important as restoring cerebral blood flow for improving long-term functional recovery after ischemic stroke.

## AUTHOR CONTRIBUTIONS

M.A., A.G.E. and A.M. conceived the project. M.A. conducted the experiments. M.A. collected and analyzed the data. A.G.E. and A.M. secured funding and provided infrastructural support. M.A. and A.M. prepared the figures. F.N.S provided technical knowledge and analytical tools. M.A and A.M. wrote the paper, with input from all authors.

## ACKNOWLEDGEMENTS

The authors thank Leyre Iglesias and Laura Aguado for technical support.

## SUPPLEMENTAL MATERIAL

Supplemental material for this article is available online.

## SUPPLEMENTARY FIGURES

**Supplementary Figure 1.**
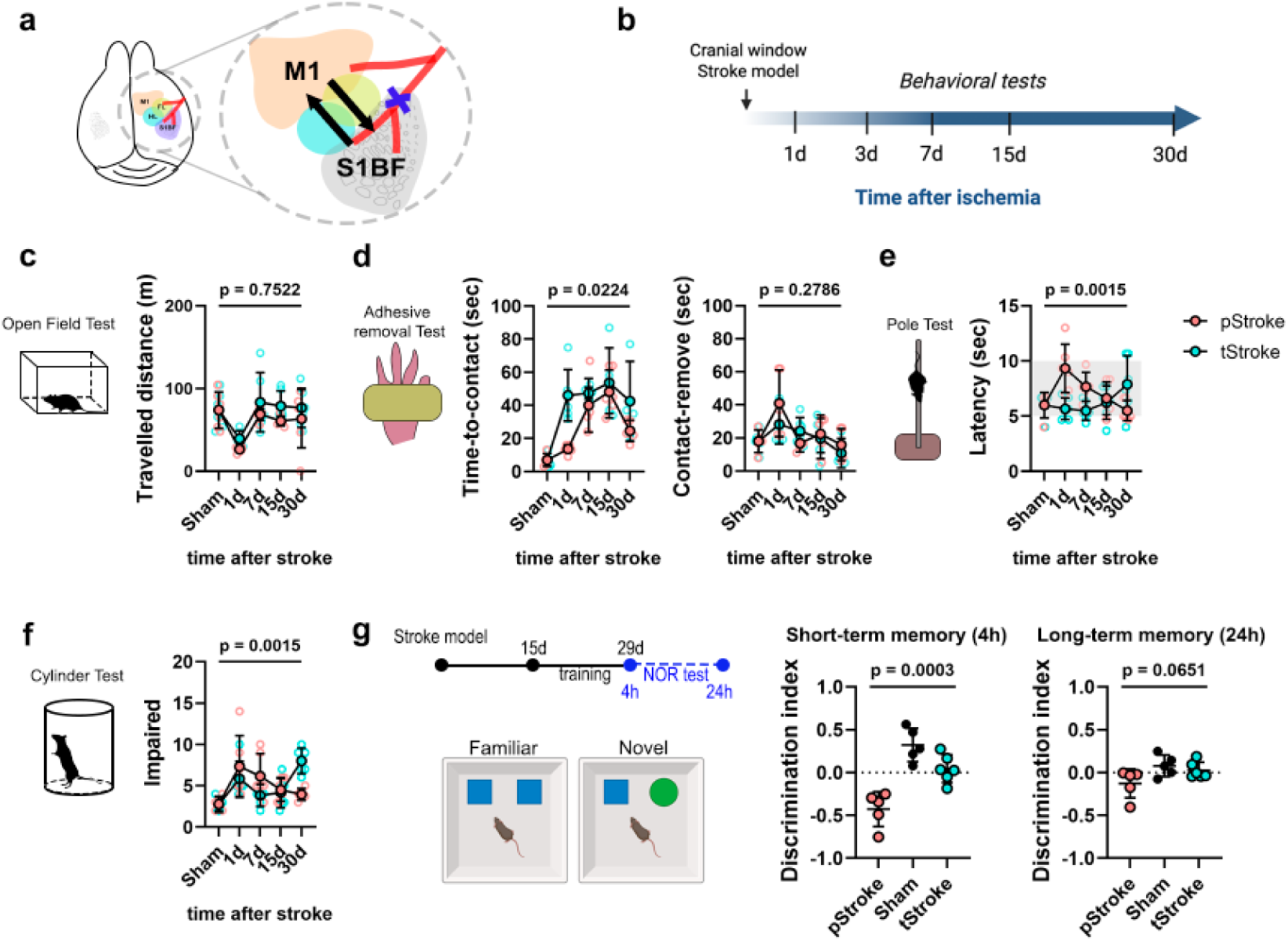
Permanent and transient stroke models produce comparable long-term behavioral outcomes. Schematic representation of the spatial relationship between the motor and somatosensory cortices (a). Experimental timeline for behavioral assessment following stroke induction (b). A battery of sensorimotor and cognitive tests was performed to evaluate functional deficits after permanent (pStroke) and transient (tStroke) ischemia. Total distance traveled in the Open Field Test did not differ between stroke groups (c). Stroke increased the latency to contact the mouth in the adhesive removal test, whereas contact and removal times were otherwise comparable between pStroke and tStroke mice, suggesting predominantly sensory rather than motor deficits (d). Performance in the Pole Test was similar between stroke groups despite differences in temporal profiles (e). Likewise, the Cylinder Test revealed no significant differences in forelimb use asymmetry between pStroke and tStroke mice (f). Recognition memory was evaluated using the Novel Object Recognition Test, and the discrimination index was determined 4 h and 24 h after the familiarization phase (g). Data are presented as scatter dot plots showing individual animals with mean ± SD.

**Supplementary Figure 2.**
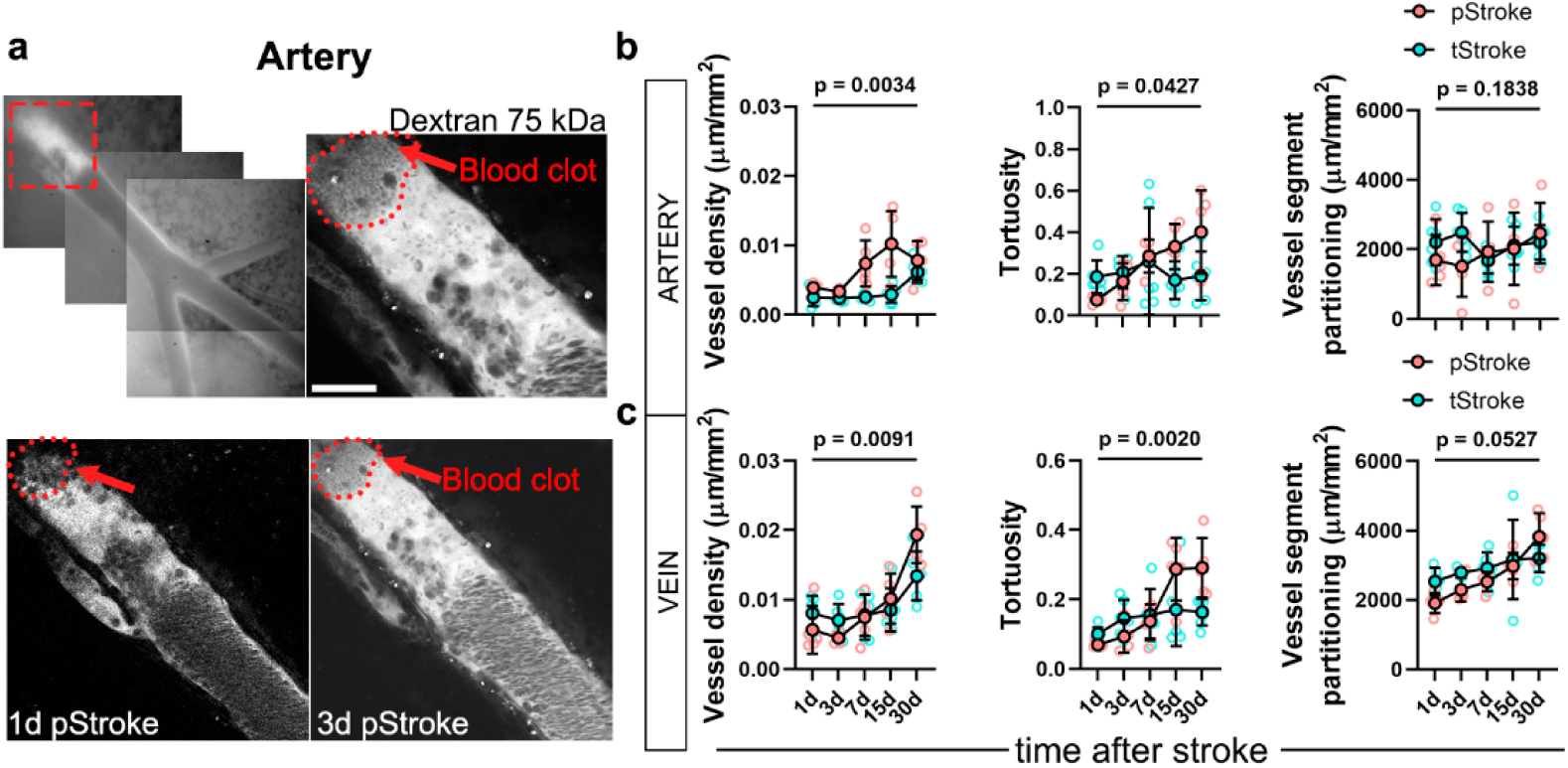
Permanent and transient ischemia induce distinct patterns of long-term vascular remodeling. Representative composite images of the ligated artery at 1 and 3 days after permanent ischemia showing blood clot formation at 3 days in pStroke mice (a). Quantification of vessel length density, tortuosity and vessel segment partitioning in the ligated artery of pStroke and tStroke mice at the indicated time points (b). Vessel length density and tortuosity exhibited distinct temporal profiles between stroke models, whereas vessel segment partitioning remained unchanged. Quantification of vessel length density, tortuosity and vessel segment partitioning in the herniated vein of pStroke and tStroke mice (c). Similar to the ligated artery, vessel length density and tortuosity differed between stroke models, whereas vessel segment partitioning was not significantly altered. Scale bar = 100 µm (a). Data are presented as scatter dot plots showing individual animals with mean ± SD.

**Supplementary Figure 3.**
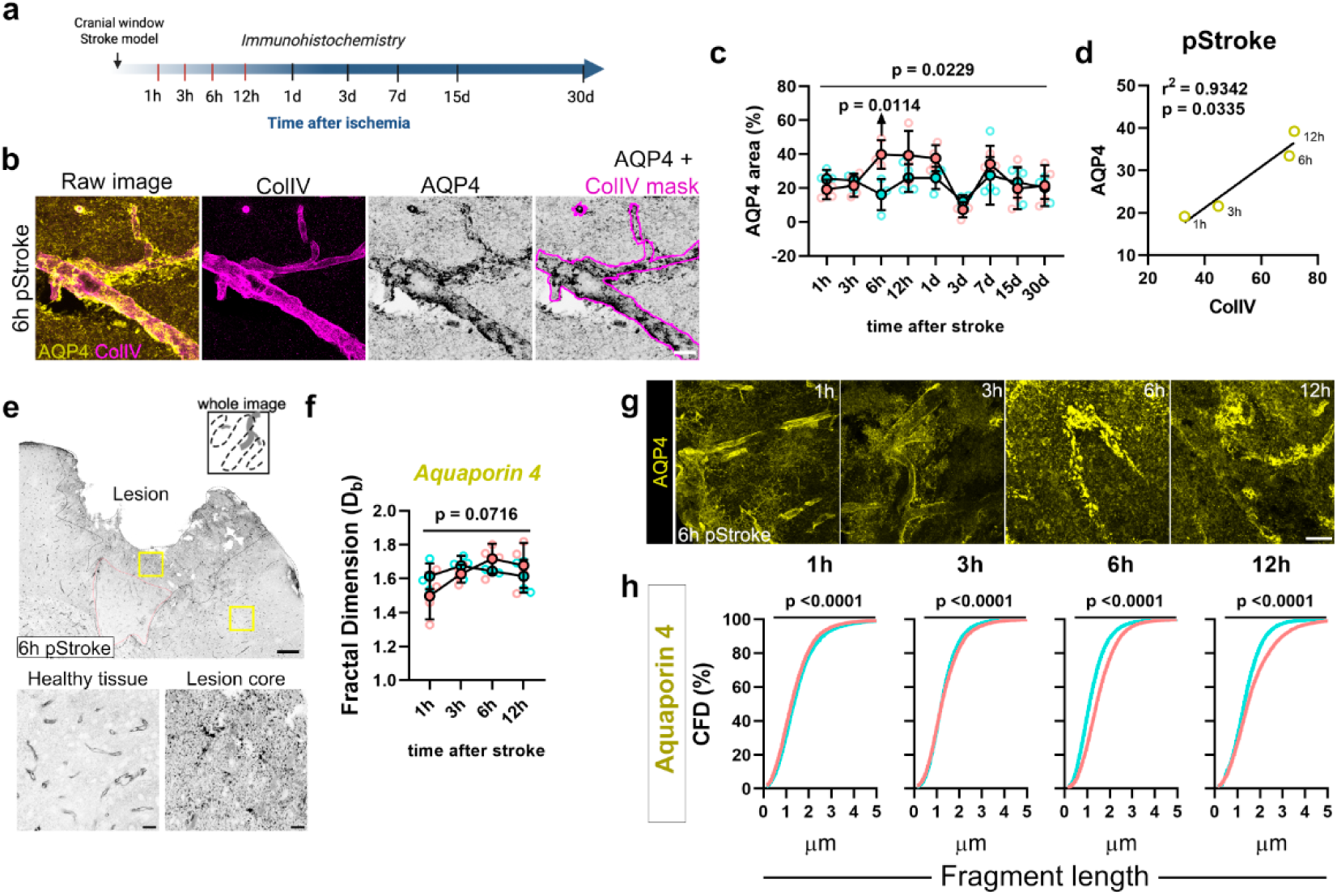
Coordinated collagen IV and aquaporin-4 remodeling during the early neurovascular response to ischemic stroke. Experimental timeline for the analysis of collagen IV (ColIV) and aquaporin-4 (AQP4) expression following permanent (pStroke) and transient (tStroke) ischemia (a). Representative immunofluorescence images showing the colocalization of ColIV and AQP4 at 6 h after pStroke (b). Quantification of AQP4 expression revealed a transient increase at 6 h in pStroke mice compared with tStroke mice (c). Correlation analysis demonstrated a significant association between ColIV and AQP4 expression during the acute phase following pStroke (d). Representative low-magnification image of AQP4 immunofluorescence in the ischemic cortex of a pStroke mouse at 6 h (e). Fractal dimension analysis revealed no significant differences in AQP4 structural complexity between stroke models (f). Representative images illustrating the temporal pattern of AQP4 expression following pStroke and tStroke (g). Quantification of AQP4 fragment size revealed significant differences between stroke models, indicating enhanced AQP4 aggregation around vessels with high ColIV expression after permanent ischemia (h). Scale bars = 20 µm (b), 200 µm and 20 µm (e), and 20 µm (g). Data are presented as scatter dot plots showing individual animals with mean ± SD.

